# PRECISE TBI Model Catalog: Increasing Accessibility and Reproducibility in TBI research

**DOI:** 10.64898/2026.02.10.704562

**Authors:** Monique C. Surles-Zeigler, Lauren Holmes, Troy Sincomb, Maryann E. Martone, Jeffrey S. Grethe, Adam R. Ferguson, C Edward Dixon, PRECISE-TBI corporate authors

## Abstract

Preclinical traumatic brain injury (TBI) research relies on experimental models that vary by mechanism, parameters, surgical procedures, species, strains, and ages, to name a few. While these models are crucial for understanding injury mechanisms and testing therapies, the progress in translating this knowledge to the clinic has been limited. This is in part due to fragmented resources and inconsistent reporting of critical variables. Here, we introduce the PRECISE-TBI model catalog, a centralized, queryable resource that consolidates metadata from published studies. The catalog integrates curated annotations from more than 450 papers, including details such as age, sex, strain, model type, device, and injury parameters. Where available, entries are also linked to protocols and datasets to enhance transparency and reproducibility. The Model Catalog serves as a living resource that enables cross-study comparison, identifies gaps in reporting, and connects the literature to datasets, protocols, device information, and other relevant resources. Analysis of the initial catalog entries revealed gaps in the reporting of device, age, and weight. In contrast, the reporting of sex improved over time, with over 90% of recent studies within the catalog papers reporting sex. Strain was also reported in most studies, with consistent reporting of specificity, especially for the C57 mice substrain. We expect the Model Catalog to serve as a valuable tool to enhance study design and reproducibility in preclinical TBI research while advancing FAIR data principles in the TBI field.

## Introduction

Preclinical models of traumatic brain injury (TBI) have been developed and used in multiple species to understand the mechanisms in humans for over a century ^1–4^. These models have been designed to study injury biomechanics, discover pathological mechanisms, and develop and test therapeutic interventions to reduce TBI-induced morbidities. Substantial additions to and improvements in available injury induction methods have occurred over the past several decades, including expanding injury induction methods, enhancing model consistency and reproducibility, and improving control over confounding variables ^1,5–15^. In addition, several new models have been recently developed to reproduce human concussion and military-relevant TBI, especially to model repetitive injury events.

Despite the multitude and continued refinement of preclinical TBI models, no therapy, devised and tested experimentally, has yet been shown to provide consistent, clinically meaningful, long-term outcomes in human TBI patients ^16,17^. Records from studies that implement these models, such as notes, protocols, and data, are scattered across several places, including but not limited to journal articles, book chapters, repositories, computer hard drives, and lab notebooks. This makes it very difficult to determine the fidelity of the models used and the associated parameters required to induce injury reproducibly. To find this information, each investigator must independently track it, including conducting multiple literature searches to locate relevant papers and their available datasets, and gathering information from various sources, such as techniques learned by visiting other labs.

Several recent efforts have been made to centralize numerous resources, including repositories to identify study protocols (protocols.io^18^), TBI-specific datasets in the Open Data Commons for TBI (ODC-TBI^19^), and Research Resource Identifiers (RRID)^20^, to increase the transparency of research metadata and enhance reproducibility. In addition, groups such as the PREClinical Interagency reSearch resourcE-TBI (PRECISE-TBI, RRID:SCR_022252) have been created to “elevate rigor, reproducibility, transparency, and translation in TBI research” by integrating and harmonizing preclinical resources and metadata, and by discovering new ways to enforce the FAIR principles in TBI research.

Here, we introduce the PRECISE-TBI Preclinical Model Catalog (Model Catalog, RRID:SCR_024626), an online queryable resource that allows users to search the metadata of preclinical TBI papers, including age, sex, weight, device, outcome assessments, TBI model, and injury parameters. In addition, we assess the completeness, consistency, and interpretability of a subset of reported preclinical TBI metadata within the Model Catalog. Additionally, it provides access to other preclinical TBI resources through the preclinical catalog toolbox, which includes protocols, a bibliography of preclinical papers, and TBI device vendors, thereby establishing a digital hub for preclinical TBI research. The model catalog currently contains over 400 curated entries, comprising a diverse range of models. This paper describes the initial phase of Model Catalog development, including the curation workflow and infrastructure. It presents quantitative analyses of the associated papers within the Model Catalog, including inter-curator agreement, metadata completeness, and reporting trends over time. These analyses establish a baseline for understanding how preclinical TBI models are currently reported and where standardization efforts could have the greatest impact.

## Methods

Here, in this section, we outline the high-level process of identifying papers for inclusion into the catalog, annotating and extracting details about these studies, and adding them to the SciCrunch Infrastructure to be viewed on the Model Catalog webpage (https://scicrunch.org/precise-tbi/about/model-catalog).

### Paper Inclusion Criteria

To create the PRECISE-TBI model catalog of preclinical TBI models, we have implemented a workflow to identify and incorporate relevant research. Our primary focus was on four TBI models: controlled cortical impact (CCI), fluid percussion injury (FPI), blast injury (BI), and weight-drop injury (WDI), as these are the most commonly used models in preclinical TBI research. However, our ultimate goal is to create an inclusive catalog that encompasses all published TBI models.

A PubMed search using specific search queries (Table 1) was used to identify articles related to the focused models. This query was run quarterly to capture new journal articles. Review papers were excluded from the Model Catalog. Once identified and curated, these new articles are added to the bibliography section of the Model Catalog website (https://scicrunch.org/precise-tbi/about/Recommended%20bibliography). Next, they were queried through a detailed annotation process that involved extracting key information and categorizing the studies based on various parameters. The annotated information was then integrated into the main model catalog, providing researchers with a searchable database of preclinical TBI models.

**Table 1:** PubMed Search Terms for Preclinical Models.

| Model | Search terms |
| --- | --- |
| WDI | "Weight drop" and TBI<br>"Weight drop" and concussion<br>"Marmarou model"<br>"Closed head" impact.<br>"Impact acceleration"<br>"concussion and Rat or mouse" |
| FPI | "fluid percussion"<br>"fluid percussion" and TBI<br>"fluid pulse" and brain |
| CCI | "Controlled cortical impact"<br>"Controlled impact"<br>"Rigid indenter" and "Brain"<br>"Pneumatic impactor"<br>"Electromagnetic impactor" and brain<br>"Leica One" Brain |
| Blast | Experimental blast<br>Open-field blast<br>Blast and shock tube<br>Blast and "advanced blast system"<br>Military and experimental TBI |

**Table 2:** Percent Annotations per Match Category. Strict agreement indicates exact matches between the values extracted for a tag, while soft agreement counts near-equivalent values as matches (e.g., synonymous terms or minor formatting differences).

| Tag | Annotation |  |
| --- | --- | --- |
|  | Strict Agreement % | Soft Agreement % |
| *Strain* | 87.77 | 88.49 |
| *TBI Model* | 71.67% | 95.1% |
| *Sex* | 93.48 | 93.48 |
| *Species* | 90.39 | 91.66 |
| *Age (Weight)* | 94.96 | 94.96 |
| *Age (Weeks)* | 71.74 | 71.74 |

**Table 3:** Intercurator Agreement per Match Category. Strict agreement represents exact matches between curators’ annotations; soft agreement counts near-equivalent values as matching. Kappa indicates agreement beyond chance.

| Intercurator agreement - metadata tag | Strict Agreement % | Soft Agreement % | Kappa Score |
| --- | --- | --- | --- |
| *Strain* | 88.9 | 88.9 | 0.88 |
| *TBI Model* | 76.3 | 95.1 | 0.82 |
| *Sex* | 93.4 | 93.4 | 0.788 |
| *Species* | 89.5 | 91.4 | 0.843 |
| *Age (Weight)* | 71.7 | 71.7 | 0.942 |
| *Age (Weeks)* | 94.3 | 94.3 | 0.707 |

## Annotations

Hypothes.is (RRID:SCR_000430)^21^, a web-based annotation tool, was used to annotate the Methods sections of the identified TBI papers. Relevant metadata about the study design (e.g., species, strain, sex, weight, age) and TBI-specific metadata (e.g., TBI model, TBI device, TBI device manufacturer, assessments, and injury specifics) were annotated. At least two curators annotated all papers. To maintain clear distinctions between the annotations made by each curator, each was assigned to a separate Hypothes.is group. Annotations were made for each paper to record identifying information, including PMID (PubMed ID), DOI (Digital Object Identifier), authors, and title.

Annotations were pushed weekly to a Google Spreadsheet for review and to identify any apparent mislabeling. In addition to the Hypothes.is. The API (application programming interface), Pandas (RRID:SCR_018214), and Gspread (RRID:SCR_027866) Python packages were used to extract and organize annotations for further processing. The workflow ensured accuracy and accessibility of the annotations.

### Inter-annotator agreement

To assess the consistency of the annotation process performed by multiple curators, two statistical measures were used: percent agreement and Cohen’s kappa. The percent agreement measures the proportion of instances in which curators assigned identical tags to a given item. This was calculated for each annotation tag and each alignment category. To better understand inter-rater reliability, particularly accounting for chance agreement, Cohen’s kappa was also computed. For the kappa statistic, the overall percent agreement was further broken down into singular, one-to-one annotations to ensure accurate calculation.

Cohen’s kappa (κ) is calculated using the formula:

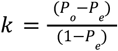

To determine Cohen’s kappa, Po represents the agreement value between curators, which is the percentage of times the curators agree on a particular annotation, and Pe is the probability that both curators would randomly agree on an annotation independently of each other.

### Annotation Alignment

Annotations were aligned and manually checked to ensure consistency and to calculate the inter-annotator agreement between the curators. This manual check also served as a pilot study, informing the development of a subsequent programmatic alignment process. During this review, the PMIDs served as the basis for alignment, enabling easy comparison between curators, although these annotations were not available for all papers. A uniform resource identifier (URI) was present for all annotations through Hypothes.is metadata. Therefore, the first significant step was to assign a corresponding PMID to each URI. This was achieved by cross-referencing the Hypothes.is metadata associated with the source documents, and extracting the PMID for each URI.

Once all URIs (and their associated annotations) had a PMID, all duplicate PMIDs for a specific curator were merged. Curators often generated multiple URIs for the same paper because annotations were created during different sessions or on various computers. The merging occurred either by combining them as comma-separated annotations in the same row or by attaching new annotations to an existing curator’s URI. This step consolidates annotations from various curators for each PMID into a single entry.

Next, the annotations were displayed side by side to categorize them across curators. For this step, we also accounted for synonyms, case differences, and white-space differences. The alignments were classified into two broad classes: “Match” and “No Match”. Annotations were assigned as a “Match” when both curators had exact or synonymous matches. For instance, if CCI and Controlled Cortical Impact were identified as synonyms. A “No Match” was recorded when two curators annotated the same PMID with differing or dissimilar interpretations. The insights gained from this alignment process, combined with inter-annotator agreement calculations, will help enhance the consistency of subsequent annotations.

### Model Catalog Infrastructure

After the Model Catalog metadata was aligned, it was added to a Google Sheet. From the Google spreadsheet, the data were ingested and transformed for addition to the Model Catalog. The Model Catalog uses Foundry, a scalable ETL (Extract, Transform, Load) platform, to extract, standardize, and integrate data. The metadata from the Google Sheet was then added to the Foundry GitHub repository to generate sample JSON records ^22^. Once validated, transformed records were reviewed and committed. After successful quality control (QC), data ingestion was finalized by registering the source into Foundry using a Command Line Interface (CLI)-based management tool. Ingested and transformed records are indexed via Elasticsearch and made available for public search through the SciCrunch Model Catalog interface (https://scicrunch.org/precise-tbi/about/model-catalog).

## Results

### Intercurator agreement

An intercurator agreement analysis was conducted to assess consistency across multiple metadata fields among curators for articles in the model catalog. The agreement was evaluated using both strict matching and soft matching (accommodated text normalization, e.g., case-insensitive matching, whitespace trimming, and recognition of curated synonym sets, e.g., “male” and “m” treated as equivalent). There was a high level of agreement (>90%) in both strict and soft matching for age (in weeks), sex, and species, indicating that, despite some variations in formatting and terminology, curators mostly agreed on these fields.

### Model Catalog Statistics and Lessons learned

As discussed in the Methods section, the terms in Table 1 are run quarterly and added to the bibliography. Currently, the bibliography includes 431 weight drop, 1,880 controlled cortical impact, 53 blast, and 56 fluid percussion papers. These papers are added to the curation queue for inclusion in the Model Catalog. To better understand preclinical TBI research papers, we examined the existing cohort in the model catalog. As of June 2025, the model catalog contains 498 records, covering 488 unique papers. Since each record in the model catalog describes metadata for one preclinical model, five papers have two entries as they represent multiple preclinical models. The model catalog currently contains the following preclinical model records: 366 weight drop, 71 controlled cortical impact, 26 blast, 15 fluid percussion, and 12 models labeled as ‘other’. Models classified as ‘other’ in the model catalog are not currently included in the focused model list (CCI, FPI, Blast, WD), but will be added to the focus list in future iterations. Of the 498 entries, most studies included only male participants (n = 386), followed by studies with no sex reported (n = 46), those involving both male and female participants (n = 35), and those with only female participants (n = 31). Additionally, these studies were conducted in multiple species, including rats (n = 249), mice (n = 226), pigs (n = 11), ferrets (n = 4), zebrafish (n = 4), rabbits (n = 2), and monkeys (n = 1).

#### Missing data

Despite established guidelines such as the Animal Research: Reporting of In Vivo Experiments (ARRIVE) guidelines (2010/2020)^23–26^, the NIH’s Sex as a Biological Variable (SABV) policy (2016) ^27^, and the Findable, Accessible, Interoperable, Reusable (FAIR) data principles (2016)^28,29^, challenges remain in achieving consistent reporting. To better understand the metadata level, we evaluated the completeness of paper metadata in the model catalog to identify data gaps. In particular, we examined the completeness of reporting age, weight, model, species, and device details, as these details significantly impede reproducibility (Fig. 1). Of all the fields examined, the device name (9%) was reported the least often, as many authors did not provide the specific device name used, but rather the type of device, such as a controlled cortical impactor rather than the name of the impactor, along with the manufacturer’s name. This was followed by age (44%) and weight (69%). Regarding age and weight parameters, most studies (44%) used weight alone, while only 19% of the records reported age alone. In comparison, 25% reported both age and weight. Sex (9%), strain (5%), species (.2%), and TBI model (100%) were the most complete.

**Figure 1.**
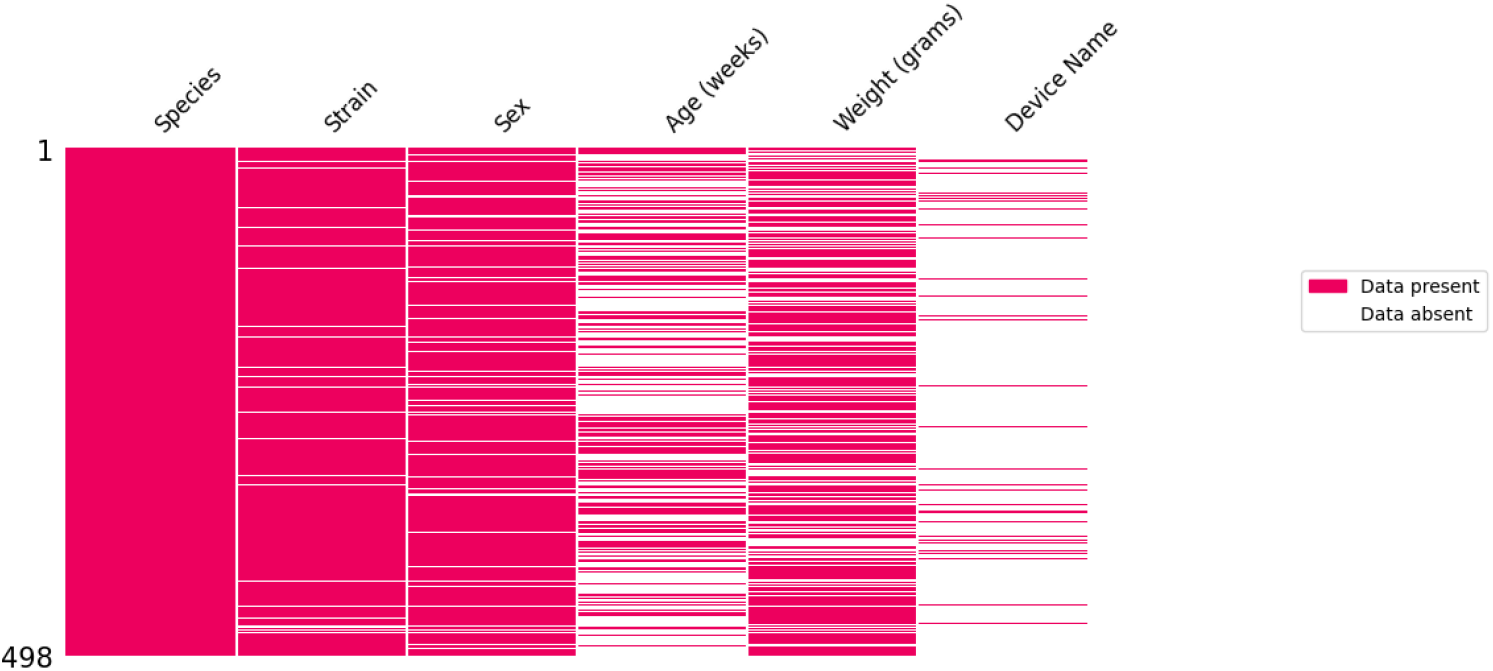
Missing data analysis. A missing data plot was created to visualize patterns of missingness in the metadata, where the rows represent papers, the columns are extracted metadata text, and a summary of the model and sex counts is included.

We then examined the reporting of these variables, organized by publication year, to assess the completeness of reporting across studies. Reporting of sex demonstrated the most significant and consistent improvement after the introduction of key reproducibility policies, notably the NIH SABV policy in 2016 (Fig.2). Before 2016, most studies were performed on males, perhaps leading to a lack of explicit reporting of sex, resulting in high missingness rates across models such as Blast and WD. While there is some year-to-year variation in reporting, a noticeable increase has been observed in the inclusion of sex data and in the reporting of both sexes in some studies. Low completeness rates have not returned to pre-policy levels, especially within CCI models. This aligns with the SABV policy becoming a funding requirement, which is likely to influence authors’ behavior. Nonetheless, less standardized model types still frequently lacked sex details.

**Figure 2.**
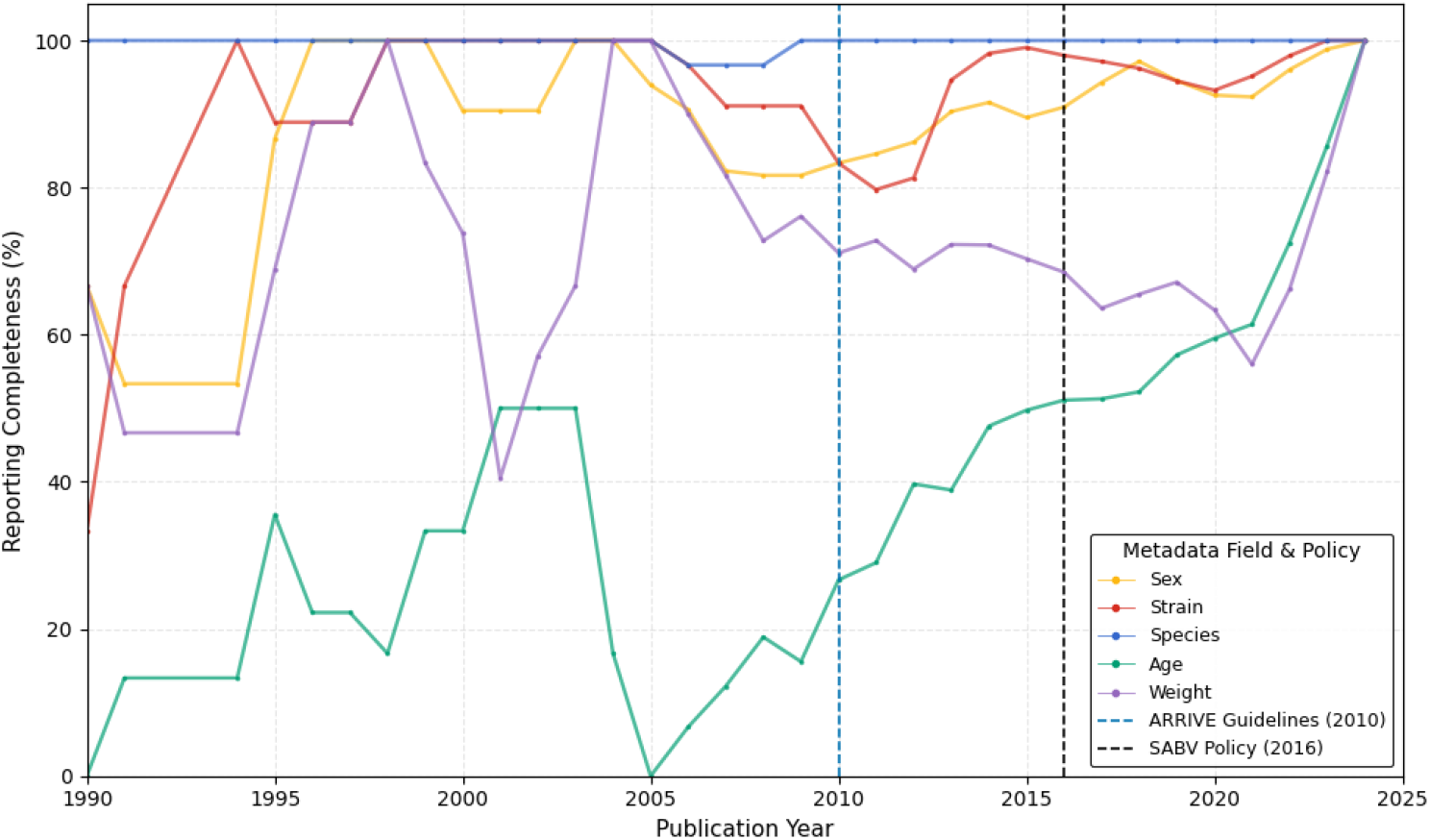
Reported Model Catalog TBI metadata over time - missing data over time. An illustration of the percentage of metadata for sex, strain, age, and weight across the preclinical TBI papers in the model catalog. Each line represents the proportion of missing values in that publication year.

Strain reporting showed steady improvement in completeness over time, similar to sex. By 2015, strain had been reported in nearly all studies, although the associated substrain remained inconsistent. A key example is the reporting of the C57 mice substrain, for which many studies did not clearly document the exact substrain used. This omission is particularly problematic because accumulated genetic drift leads to distinct variation over time, making the substrain used vital for studying reproducibility across studies. For instance, there are known differences between C57BL/6J and C57BL/6N, including the metabolic response^30–32^, behavior^33–35^, and seizure susceptibility^36–38^. Of the C57 strain reported, only 47% reported whether it was C57BL/6J (n=55) or C57BL/6N (n=7). Even within these reports, there were variants of C57BL/6J reported c57bl/6j (n=43), c57bl6/j (n=8), c57b6/j (n=4).

Age and weight are also critical pieces of metadata that can increase the reproducibility of studies, but these have been consistently underreported. There was some improvement in the reporting of age, which increased from 16% before 2010 to about 56.2% after 2016 (Table 4). These trends were species-dependent, with age mostly reported in mouse studies, weight mostly reported in rat studies, and both reported inconsistently in larger species (e.g., pigs and monkeys). Although a few CCI studies in this cohort (n=2) published after 2017 began reporting age and weight, these were exceptions. Models like Blast and WD often omit these fields or use vague descriptors (e.g., adult and young) that are helpful when combined with age and weight but are not useful on their own for reproducibility. Unlike sex and strain, which showed policy-driven improvements, there was no strong trend toward increased reporting of age or weight following the ARRIVE or SABV guidelines. This suggests that while reproducibility guidelines have improved the reporting of sex, additional efforts are needed for the reporting of age and weight.

**Table 4:**
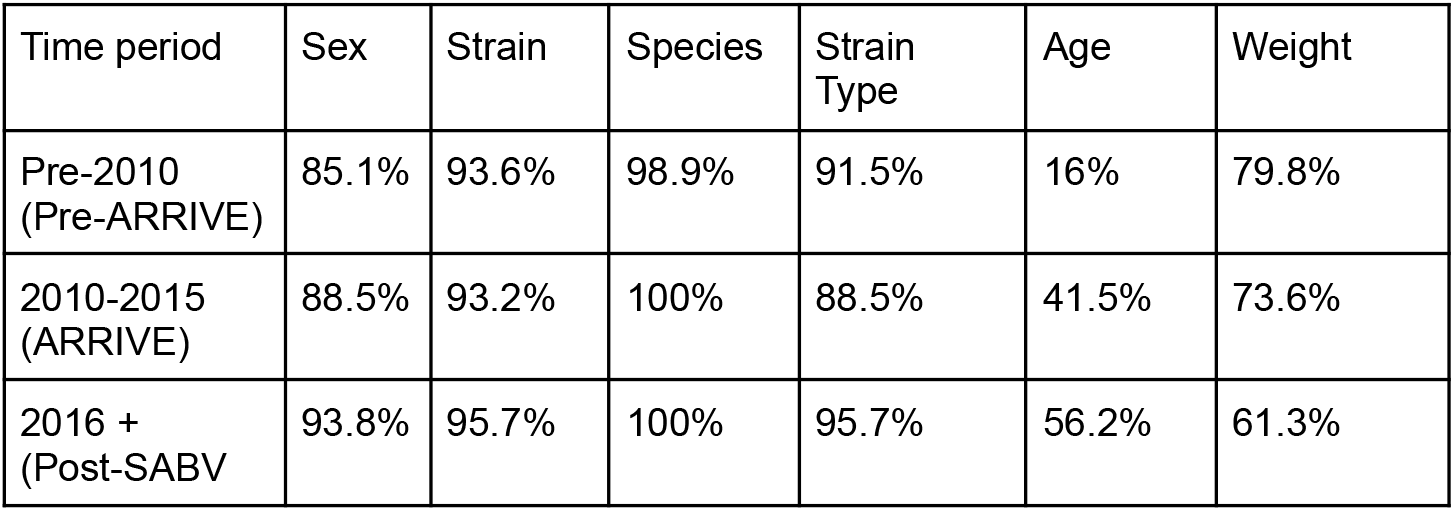
Completeness of metadata in papers by policy period.

#### Inconsistent reporting of model names

The reporting of preclinical TBI model names varied significantly in terminology. In many cases, the same underlying model was described using different names, spellings, or descriptors, which complicated the cataloging and comparisons across studies. For example, FPI reporting was consistent, with standard terms such as FPI, lateral FPI, and midline FPI. In contrast, there were also non-specific variants reported in the CCI model, such as terms like Modified CCI or closed head CCI, which blurred the distinction between the defined injury or instrument used and the model itself. Traditionally, a classical CCI typically refers to an open head injury, so the addition of ‘closed head’ or ‘modified’ refers to procedural differences that are not always captured by the name alone. This does not even include studies that do not include these descriptors but are not classical CCI (e.g., closed-head impact).

Blast and WD had even more variation (Fig. 3). Blast model reporting was consistent with recognizable named models, such as the Missouri open-field blast model, the Open-field blast, or repeated blast overpressure. In contrast, the weight drop model showed the greatest variation in its reporting, with more than 15 distinct naming variants, even when accounting for simple synonym differences (e.g., injury versus method, or hyphenated versus unhyphenated terms). Beyond the standard references to models such as Feeney’s and Marmarou’s, modified models were also reported, including the closed weight-drop model, the weight-drop contusion model, the focal weight-drop model, the weight-drop impact-acceleration method, and the concussive weight-drop model. These naming inconsistencies highlight the lack of standardization in the field, making it difficult to compare models across studies solely on names and underscoring the need for an ontology-based approach that captures shared features among models.

**Figure 3.**
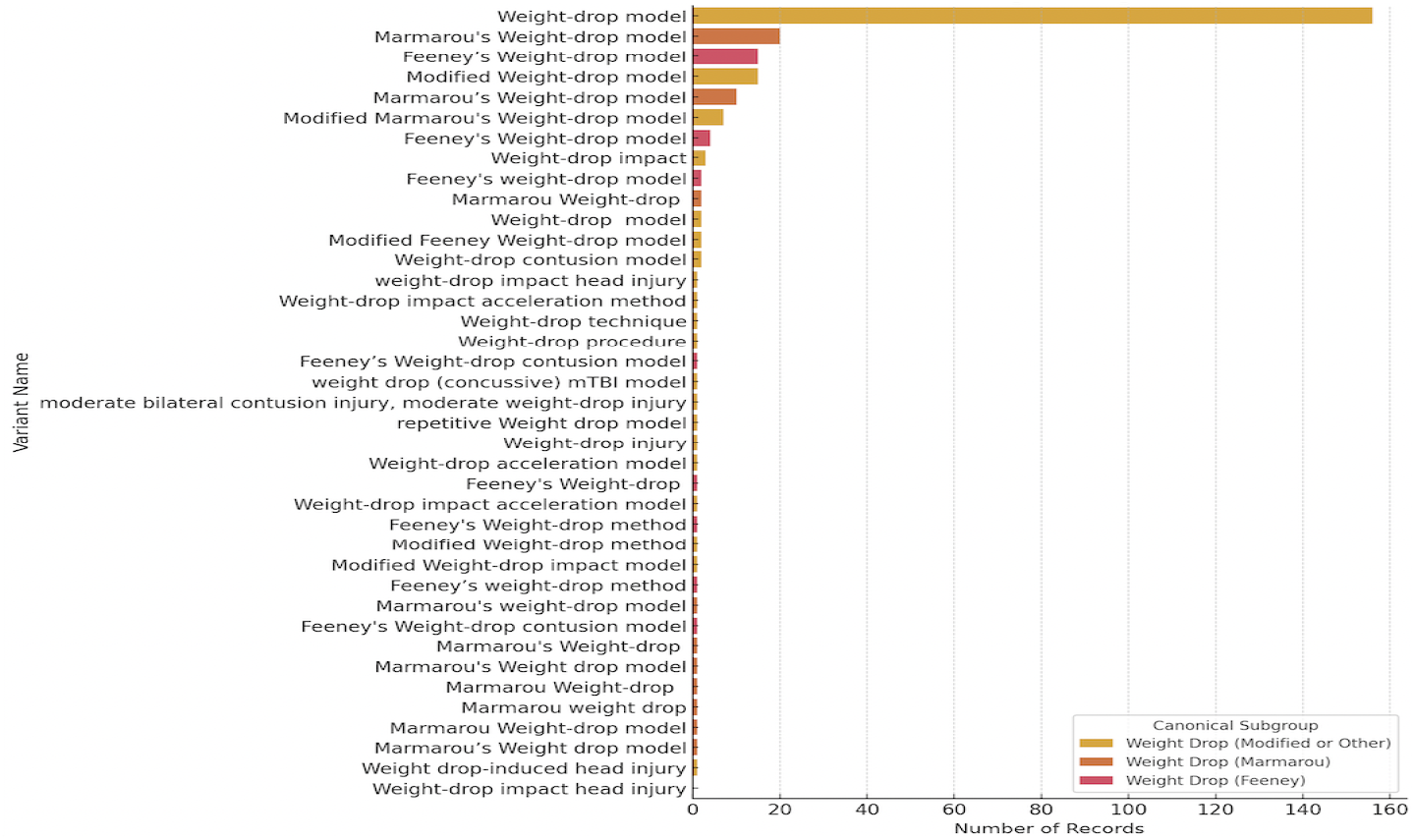
Weight Drop Model naming variation in the model catalog. The bar chart illustrates the variation in naming conventions for drop models in the model catalog. The X-axis shows the number of records in the catalog by name variant (Y-axis). Most studies reported the “Weight-drop model,” but the reporting varied across most studies.

**Figure 4.**
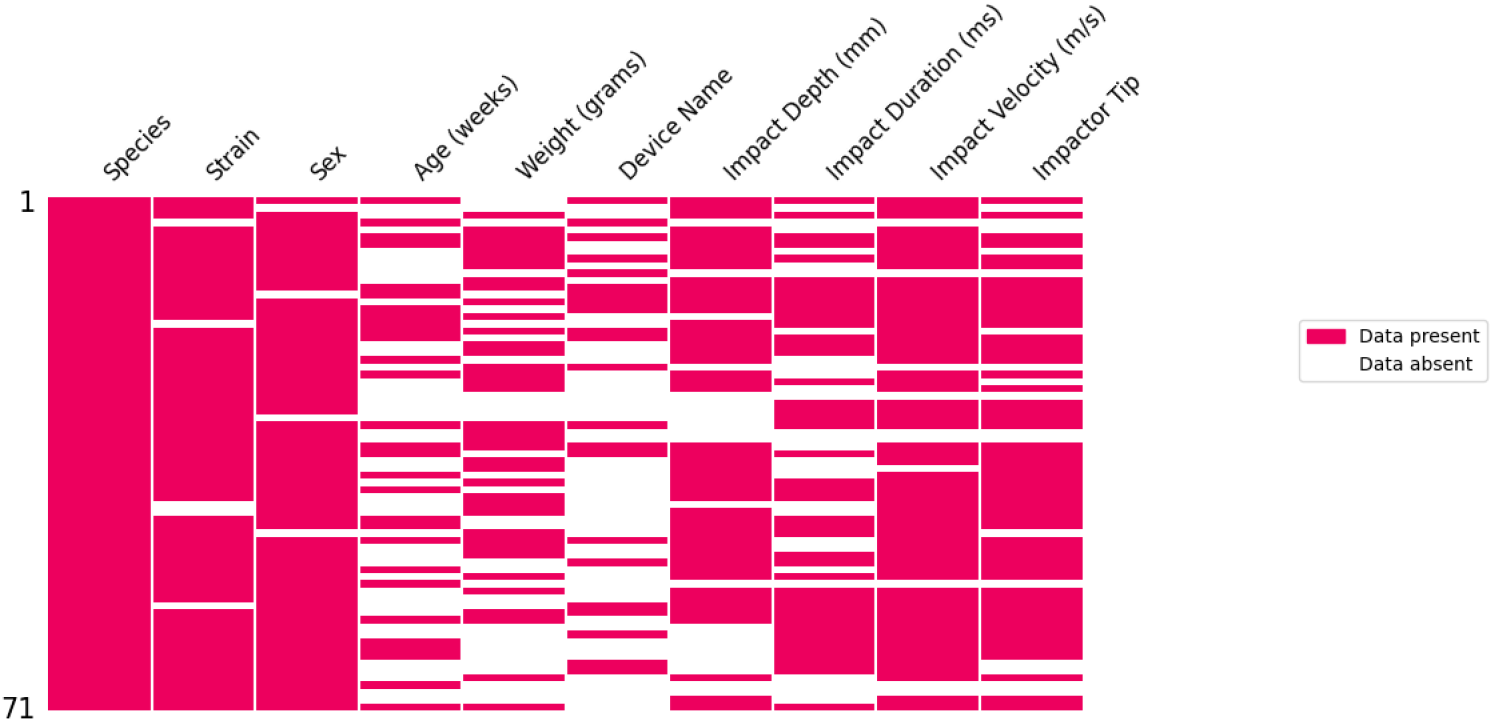
CCI missing data analysis. This illustrates the percentage of missing metadata for CCI-related papers in the model catalog, along with associated impact parameters and device information.

#### CCI missing data

Further examination of the metadata associated with the CCI model, including the impact parameters. Impact parameters related to the other models will be added in the next iteration of the catalog. Of the 71 CCI records in the model catalog, 69 records (97%) had at least one missing value. Species were reported in all entries, followed by sex (94%), strain (93%), impact velocity (86%), impact tip (76%), and impact depth (69%). As in previous analyses of the complete model catalog cohort, weight (53%), age (41%), and device name (31%) were the least frequent characteristics. There were only 11 entries (15%) with both weight and age, while 15 entries (21%) were missing both.

To assess the relationships between key numeric variables in the CCI model entries, we used non-parametric correlation methods (Spearman and Kendall). These methods were chosen because the data are non-normally distributed, skewed, and contain outliers.

#### CCI Impact parameters reporting

We examined reporting of impact parameters within the CCI model, focusing on velocity, depth, duration, and tip shape. There is a moderate positive correlation (r = 0.52) between weight and impact depth, indicating that larger animals tend to use increased impacts. For example, monkeys and pigs had greater impact depths (∼5-7 mm), whereas rodents had shallower depths, which may be related to differences in cortical mantle thickness (Koo et al., 2012; Hutsler et al.,^39,40^. Rodents had the highest impact velocity (4.5 m/s) when compared to other species, with pigs and monkeys having an average of 3-4 m/s. Regarding impact duration, it varied across species, with monkeys having the longest duration (∼150 ms), followed by pigs (∼130 ms), ferrets (∼100 ms), and mice (∼74 ms). Of those reporting impact tip shape in CCI, flat impactor tips were used in rodents, while larger animals had more beveled and rounded tips. This reveals differences in these impact parameters that may independently contribute to morbidity.

### Model Catalog Walk-through

The model catalog landing page is located at https://scicrunch.org/precise-tbi/about/model-catalog (Figure 5a). The landing page provides users with a brief description of the catalog’s goals, key facts, and frequently asked questions (FAQs), along with links to the TBI model pages and the model catalog.

**Figure 5.**
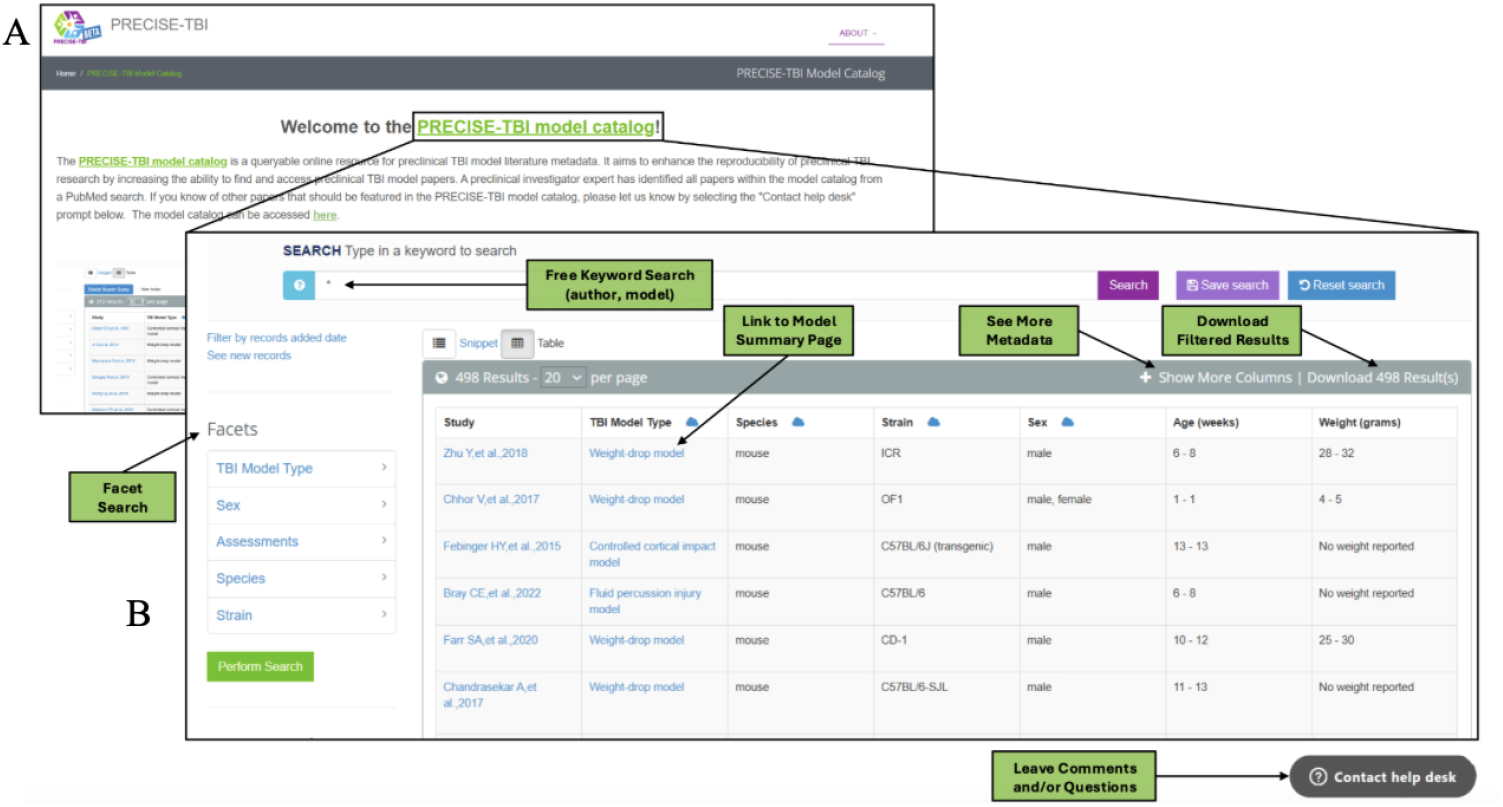
Landing page and table view of the model catalog. (a) Depicts a snapshot of the model catalog landing page accessible at https://scicrunch.org/precise-tbi/about/model-catalog. While (b) illustrated an annotated view of the PRECISE-TBI Model Catalog user interface, which highlights key features of the searchable interface for the model catalog, such as free keyword search, facet filters, and the model summary page.

The model catalog is freely accessible to all users. There is an option to register for an account on the SciCrunch website by providing a name, email address, password, and optionally, an associated institution. Once registered, this allows access to additional features in the model catalog, such as saving searches and accessing the application programming interface (API), as well as other resources within the SciCrunch Infrastructure ^20,41–45^.

#### Metadata views

When users access the model catalog, they are taken to the snippet view. In this view, records are displayed in a vertical list, with each entry shown as a block of information about a specific record (preclinical TBI paper metadata). Each block begins with a short citation as the heading, which is hyperlinked to the article metadata within the SciCrunch Infrastructure. Below the heading are key details of the preclinical study entry, including the model type, species, strain, assessments, and citation. The snippet view offers a more compact view of each search entry. The table view uses a spreadsheet-like format, with the first row as column headers and subsequent rows containing study metadata. The view provides supplementary data, publications, protocols, and related resources via embedded hyperlinks, offering further insight into the model’s research applications. Users can view additional columns in the table by selecting “Show More Columns.” This includes access to TBI Device, Device RRID, Impact parameters, TBI model, Strain, Model catalog ID, links to PubMed through the PubMed ID column, Associated Protocols in protocols.io, and Associated Datasets in ODC-TBI. This view also allows columns to be sorted by text filter or in ascending and descending order, making it easier to identify the most relevant options for a given research question. In the table view, users can save their search results and download them as a CSV file to their local computer. Additionally, users can select the cloud icon next to the column headers to display a word cloud, where the size of a term reflects its frequency in that column.

Regardless of the view, five features are present in the catalog: free-text search, preclinical TBI article metadata view, facet search (side panel), a “Contact Help Desk” button, and access to the Model Catalog Toolbox landing page. The free-text search box is located at the top center of the page. Below that are the entries of the TBI paper metadata records, which can be viewed as a snippet or in table view. The snippet and table views provide different views of the preclinical TBI metadata in the model catalog, including article citations, species, TBI models, strains, and assessments. On the left side of the screen is the faceted search. Towards the top of the model catalog page, users can select the “PRECISE-TBI” image to go to the *PRECISE-TBI toolbox landing page* or select the “Contact Help Desk” button at the lower right. The features of these pages will be discussed below.

#### Free text search

There are multiple ways to search and filter data within the model catalog. A traditional search bar is located at the top of the catalog. This search box enables you to search the metadata in the model catalog database, including author last name, model, sex, and species. If logged in to the website, users can save their searches for future use by selecting the ‘Save search’ option to the right of the search bar. Similarly, any search done on the catalog can be reset by pressing the ‘Reset search’ button.

#### Preclinical TBI article metadata view - Links to TBI Protocols and Datasets

To promote reproducibility and transparency within the TBI community, the model catalog links article metadata to other related resources within the table view. These resources, including protocols and datasets, are housed in repositories such as protocols.io and ODC-TBI. The model catalog provides links to these resources under appropriately named columns.

The PRECISE-TBI space on protocols.io enhances FAIR data standards in the TBI field by providing detailed research protocols. Protocols.io, an open-access repository, facilitates the sharing of research protocols in a way designed to foster their use. To enhance protocol accessibility and reproducibility across laboratories, this platform assigns a unique, permanent Digital Object Identifier (DOI) to each uploaded protocol. Investigators can modify these protocols by creating new versions or forking them, while always maintaining a clear link to the original. While researchers can submit protocols directly to protocols.io, PRECISE-TBI also offers a service to add protocols, with the guidance and approval of an investigator, to the PRECISE-TBI workspace. To initiate this pipeline of submitting to the PRECISE-TBI protocols.io workspace, investigators are asked to complete the form at https://scicrunch.org/precise-tbi/about/tbi-protocols. Currently, 12 protocols have been submitted from 7 laboratories, comprising 9 model protocols that range from closed-head injury models, FPI, and CCI (including polytrauma) to open-field blast, as well as three assessment protocols. Collectively, these protocols have been viewed over 4,000 times as of October 2025.

ODC-TBI also fosters reproducibility and transparency in TBI research by providing a data-sharing portal and repository for TBI experimental data, and by allowing users to share data privately with their team or collaborators (Espin et al., 2019; Chou et al.,^19,46^. Papers linked to these submitted protocols and datasets receive priority for curation and annotation in the Model Catalog, as they are considered the gold standard for ensuring reproducibility in TBI research.

#### Preclinical TBI article metadata view - Model Pages

The TBI model pages provide a comprehensive one-page summary for each TBI model within the Model Catalog. Users can access a Model Page (Fig. 6) for each model in the catalog by selecting the hyperlinked text in the “TBI Model Type” column. The information provided on the model page includes the definition and synonyms used to refer to the model, the PRECISE-TBI Common Data Elements (CDEs) associated with the model, and TBI model devices (LaPlaca, Harris et al., this issue). Additionally, this page provides information about the model, including species, assessments, citations, linked protocols, and data available through the ODC-TBI ^19^.

**Figure 6.**
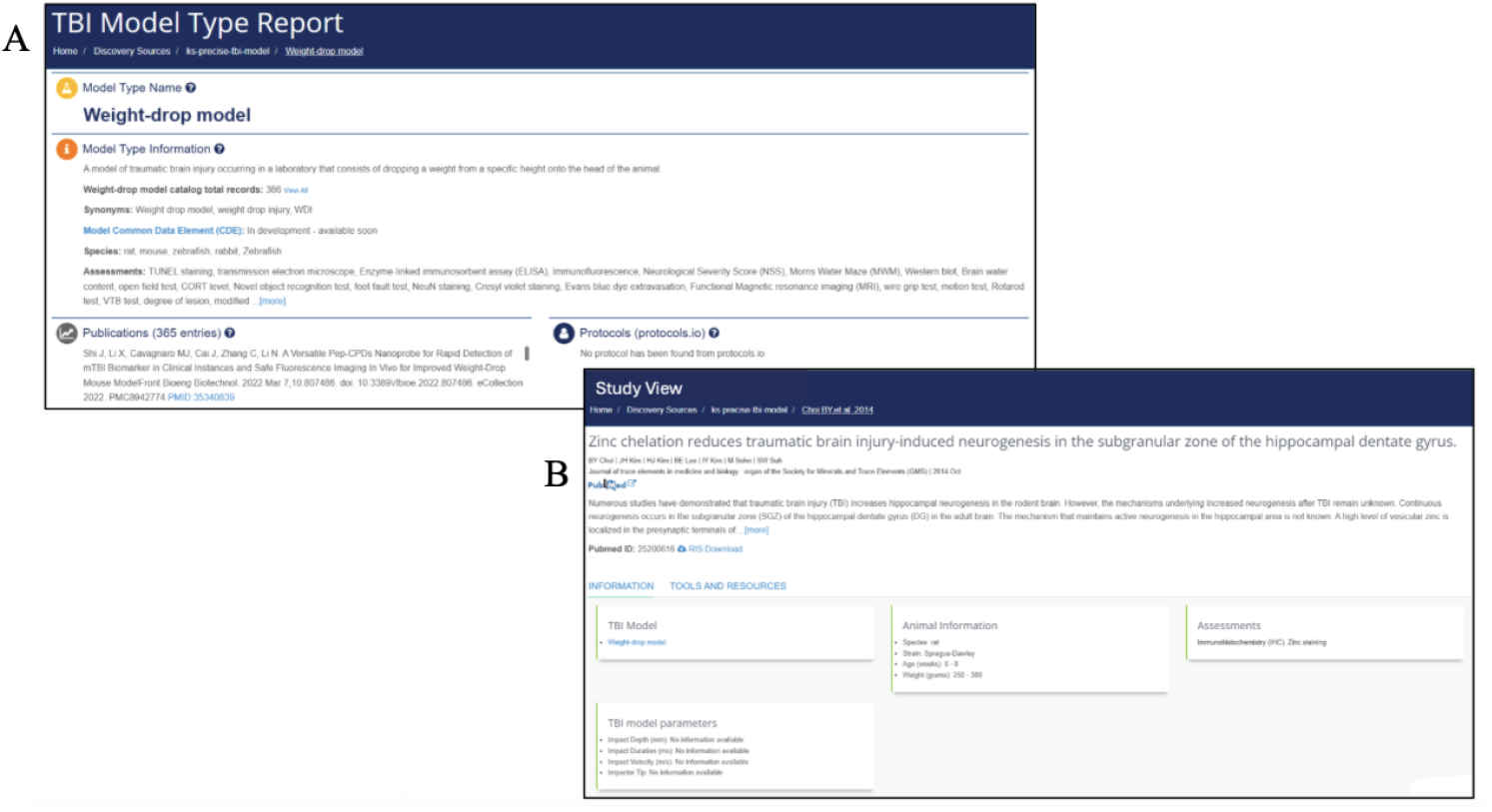
PRECISE-TBI Model Catalog’s model and study pages. (b)The TBI Model Type Report (top left) provides curated background information on a specific TBI model, including a textual description, relevant experimental considerations, associated publications, and links to protocols. (b)The Study View page (bottom right) contains metadata for an individual preclinical TBI study, including animal information (species, strain, sex, weight, and age), the assessments used, and specific TBI model parameters.

#### Faceted search

The faceted search allows users to refine their search by specific criteria, such as model type, sex, species, and strain. The search filters help users quickly retrieve information that matches their research needs. The filter can be applied to one or multiple facets. For instance, a user can select “female” under the “sex” facet or combine it with other faceted filters, such as “TBI Model type” and “Species,” to search for papers in the model catalog that focus on females of a particular species or TBI Model.

#### “Contact Help Desk” button

The “Contact Help Desk” button allows users to share feedback about the Model Catalog. The input can be related to an error, a question, a critique, or comments about the Model Catalog that the user would like to share. To complete the form, users must submit their name, email address, and a message detailing any issues or concerns.

## Discussion

Here, we present the PRECISE-TBI Model Catalog, an annotated repository of published preclinical TBI studies that systematically captures and centralizes key metadata about the animal models from multiple sources. It is a unique resource that catalogs crucial metadata about animal models, including injury type, device settings, animal characteristics (species, strain, sex, and age), and experimental designs. The catalog currently contains 498 entries, including 488 preclinical TBI papers focused on CCI, WD, Blast, and FPI models, with over 1,500 papers in queue for addition. Since the catalog is searchable and interoperable, it not only aggregates data but also provides a framework to increase reproducibility and identify reporting gaps in the field. To understand the content and coverage of the catalog, we performed exploratory data analyses across model types, impact parameters, and subject variables. The descriptive summaries revealed key patterns in these papers, providing insight into the current literature in the model catalog and uncovering several potential gaps that are not unique to the TBI field.

One substantial observation from the analysis is the lack of reporting of essential variables in preclinical TBI papers. This was most notable in variables like device name, age, and weight. Many of the articles did not provide specific device names, instead offering only generic descriptions, such as “controlled cortical impactor.” However, this is understandable in models like the weight-drop model, where the object is dropped from a variable height that may not be directly related to a standard device or available source information. This omission greatly impedes the replication of injury-induction methods. Similarly, over half (56%) of entries omitted numerical age data, and 31% lacked weight information. These missing biological variables have a fundamental influence on the response to TBI ^47–49^, thereby limiting the ability to compare results across studies to interpret the outcomes accurately. The reporting of sex increased after 2016 when the SABV policy^50^ was released, from 85.1% to 94%. It was not surprising that this improvement was not similar in the reporting of age and weight. However, age and sex reporting were emphasized in the ARRIVE guidelines released in 2010 ^23–26^, with a slight increase in age reporting from 2010 to 2016, from 40% to 56%, and a subsequent increase from 2016 onward. Weight showed the opposite trend, decreasing from 40% to 56%. These patterns suggest that policies focused on a single variable, in this case sex, may be more effective. Additionally, there is a persistent need for more substantial incentives or requirements to report age and weight.

Our analysis also identified variations in how preclinical models were reported in papers from the model catalog. Models such as the CCI models were described based on the device used to induce injury, rather than on the formal definition of a CCI model, which is an open-head impact injury ^51^. Additionally, many of these models had an unspecified descriptor, such as “modified CCI,” with no further explanation of how the model was modified. In contrast to FPI, which was reported consistently, the weight-drop model had the most variation. Even after synonym normalization, this model lacked standardized nomenclature. Analysis showed more than 15 distinct naming variants, emphasizing the need for controlled vocabularies. Also, stating that the model is modified without additional clarification in the reporting complicates comprehensive literature searches and data aggregation, making it difficult to compare analyses across studies. Actions such as adding a protocol in protocol.io are the first step. Additionally, resources such as the Brain Injury Knowledge Ontology (BIKO) can help identify ways to standardize these modifications and integrate them into a formal framework (Surles-Zeigler et al., this issue).

Another interesting finding is the incomplete and inconsistent reporting of animal strain details for C57BL/6J mice (e.g., C57BL/6J vs. C57BL/6N), particularly the omission of the exact substrain. While strain was reported in approximately 95% of entries in the Model Catalog, closer examination revealed inconsistent reporting. For example, C57BL/6J differs in metabolic response ^30–32^, behavior ^33–35^, and seizure activity ^36–38^, meaning that simply adding the label C57 without the exact substrain is not directly comparable and limits reproducibility and the ability to compare results across studies. For example, the C57BL/6J substrain has higher anxiety-like behavior, which may impact performance on specific behavioral tasks. Therefore, the exact subtrain used has profound implications for reproducibility, as it can alter the outcomes of seemingly similar studies. These types of inconsistencies may be identified during the publication review process.

Efforts to increase reproducibility and transparency in research, particularly in TBI, will help address methodological inconsistencies. Efforts such as the preclinical common data elements (CDE) aim to identify and standardize the minimal terminology and data points required for reporting across datasets or papers. While these initiatives have previously been introduced to the preclinical TBI landscape [15,46,47], recent endeavors have reintroduced and further developed these CDEs within the PRECISE-TBI project to create the PRECISE-TBI CDEs (LaPlaca, Harris et al., this issue) and integrate them with datasets in repositories such as the ODC-TBI ^19,46^. The linking of these published datasets with a DOI in ODC-TBI, with some core CDEs to the published paper in protocols.io, and the metadata in the model catalog with additional information about species and devices connected to an RRID and models, all create the necessary infrastructure for the “paper of the future”.(Gu et al., this issue) The foundation for a future digital paper, where all essential study components are interconnected and reusable, thereby increasing rigor and reproducibility in TBI research.

The PRECISE-TBI Model Catalog establishes a foundation for improving rigor and reproducibility in preclinical research by providing a web-based platform that consolidates resources, including protocols, datasets, and metadata for preclinical TBI papers. Currently, data in the model catalog has allowed us to identify consistent gaps in reporting, such as underreporting of TBI devices, age, and weight, as well as inaccurate reporting of substrains. Since the model catalog contains only a small fraction of available preclinical TBI papers, using Artificial Intelligence (AI)-assisted tools will improve metadata extraction and expand coverage of these papers in the catalog. Here, we present an analysis of the current curated paper cohort to inform the continued development of this resource. The incorporation of these additional studies will provide a more comprehensive and balanced view of the field, offering a more well-rounded understanding of reporting trends and methodological variation. These analyses also revealed patterns of missing data, indicating areas where reporting practices or curation pipelines can be enhanced. As such, future work will focus on expanding the catalog to include additional papers and TBI models, incorporating PRECISE-TBI CDEs, and adding additional links to protocols and ODC-TBI datasets to increase interoperability within the field. Together, these efforts will create a comprehensive resource that supports discovery, comparability, transparency, and reproducibility in preclinical TBI, thereby informing study design and guideline development.

